# Metabolic Strategies That Enable Oral Commensal Persistence in a Lower Airway Environment

**DOI:** 10.1101/2025.07.03.662932

**Authors:** Ashley M. Toney, Junu Koirala, Apollo Stacy

**Author notes:** These authors contributed equally (Ashley M. Toney, Junu Koirala). Author order was determined based on seniority. Corresponding author: Apollo Stacy.

## Abstract

Oral microbiota are increasingly implicated in chronic inflammatory diseases beyond the mouth, including bronchiectasis, a condition marked by persistent airway inflammation, mucus accumulation, and limited therapeutic options. Among these microbes, commensal *Neisseria*, typically considered health-associated in the upper airway, are emerging as opportunistic colonists of the inflamed lower airway. However, mechanisms supporting their persistence in this hostile environment, characterized by antibiotic pressure, nutrient limitation, and interspecies competition, remain poorly defined. Here, we show that the prevalent oral commensal *Neisseria mucosa* exhibits markedly enhanced antibiotic resistance and anoxic growth in a synthetic medium (SCFM2) that mimics sputum from individuals with bronchiectasis. Using genome-wide transposon sequencing (Tn-seq), we identified key genetic determinants of *N. mucosa* fitness, including pathways for L-lactate catabolism and pyrimidine biosynthesis. Notably, nitrate respiration was also essential for growth, linking use of this inflammation-associated electron acceptor to fitness under oxygen-limited, sputum-like conditions. Loss of nitrate reductase impaired anoxic L-lactate catabolism, reduced competitive fitness against *Pseudomonas aeruginosa*, and abrogated growth in SCFM2, as well as human saliva—a relevant, nitrate-rich nutrient source in the oral cavity. Furthermore, chemical inhibition of nitrate reductase using tungstate suppressed growth of both *N. mucosa* and *P. aeruginosa* in SCFM2, but not in standard media, revealing a context-specific metabolic vulnerability of nitrate-respiring airway pathogens. These findings suggest that inflammation-compatible traits like nitrate respiration, while supporting oral commensalism, may also drive opportunistic expansion in the inflamed airway. Targeting such pathways could offer a non-antibiotic approach to limit oral bacterial persistence in bronchiectasis and other chronic diseases.

**Importance:** Chronic respiratory diseases such as bronchiectasis and cystic fibrosis are marked by persistent infection and inflammation, with oral bacteria increasingly recognized as active contributors. Among these, *Neisseria* species, typically considered health-associated commensals, are frequently detected in the lower airway, yet little is known about how they persist in this hostile environment. Here, we identify key metabolic pathways that support the survival of *Neisseria mucosa* in sputum-like media, including nitrate respiration, a pathway closely linked to inflammation. We also show that *N. mucosa* exhibits enhanced antibiotic resistance under these conditions, underscoring the need to study microbial physiology in host-relevant contexts. Finally, we demonstrate that a selective inhibitor of nitrate respiration suppresses *N. mucosa* growth in sputum-like but not standard media, revealing a context-specific vulnerability. These findings suggest that targeting inflammation-compatible metabolic pathways may inform new, non-antibiotic approaches for managing chronic respiratory diseases.

## Introduction

The human oral cavity harbors one of the most taxonomically rich microbial communities in the body, comprising over 800 described taxa (1). Although oral microbiota are generally commensal and spatially confined to the upper aerodigestive tract, a growing body of research suggests that they can contribute to disease far beyond their primary habitat (2). In addition to local conditions such as dental caries and periodontitis, oral microbes have been implicated in a broad range of distal diseases, including cardiovascular disease, diabetes, rheumatoid arthritis, and inflammatory bowel disease. These associations are thought to involve several indirect, long-range mechanisms, including endotoxemia (3), circulation of microbially produced metabolites (4, 5), and maladaptive trained immunity (6).

Another major route of oral microbiota-driven disease is direct colonization of extra-oral sites, particularly in individuals with compromised immunity. While typically benign, oral commensals can translocate to distal niches, where they can exacerbate inflammation (7, 8) or establish opportunistic infections such as brain abscesses (9), infective endocarditis (10, 11), and respiratory infections (12, 13). This is surprising given that many oral taxa are nutritional specialists with reduced genomes and narrow ecological niches (14, 15), raising a fundamental question: how do oral-adapted microbes survive, and even thrive, in radically different environments across the human body?

One such environment is the chronically inflamed lower airway. While aspiration of oral contents has long been recognized as a cause of acute pneumonia (16), recent work has also implicated oral microbes in chronic respiratory diseases, including chronic obstructive pulmonary disease (COPD), bronchiectasis, and cystic fibrosis (CF) (17–20). While etiologically distinct, these conditions share in common persistent inflammation, impaired mucociliary clearance, and recurrent infection, which collectively remodel the airway microbiota (21). In CF and non-CF bronchiectasis, both of which are marked by irreversible airway dilation and mucus accumulation, oral taxa such as *Streptococcus*, *Prevotella*, and *Veillonella* are frequently detected in sputum and bronchoalveolar lavage samples (12, 22, 23). Though partly attributable to upper-airway contamination (24), mounting evidence points toward a functional role for these taxa in shaping community structure, immune responses, and disease severity, similar to the contributions of canonical pathogens like *Pseudomonas aeruginosa* (12, 25–28).

Among oral taxa linked to bronchiectasis, commensal *Neisseria* are of particular interest. This complex genus of Gram-negative, aerobic diplococci is notorious for the non-oral pathogens *N. gonorrhoeae* and *N. meningitidis*, causative agents of gonorrhea and meningitis, respectively (29). At the same time, the genus comprises several distinct commensal species, including *N. cinerea*, *N. elongata*, *N. lactamica*, *N. mucosa*, *N. sicca*, and *N. subflava* (30, 31). Traditionally regarded as health-associated members of the upper airway microbiota, these taxa contribute to microbial community stability (32, 33) and can modulate host immune responses (34, 35). Within the oral cavity, they can exhibit biogeographic niche specialization, with some species showing preference for specific habitats such as the tongue dorsum or dental plaque (14, 36). Notably, these spatial distributions are reflected in habitat-specific functional adaptations, with plaque-associated *Neisseria* (e.g., *N. mucosa*) showing enhanced capability for nitrogen metabolism (14).

However, recent studies have reported *Neisseria* enrichment in patients with chronic lung diseases, particularly bronchiectasis (12, 13, 21–23, 37), raising questions about their potential role in disease progression. While not considered primary pathogens, the fact that commensal *Neisseria* a) can persist in inflamed airways, b) exacerbate pulmonary disease in animal models and c) share key metabolic and virulence-related traits with pathogenic *Neisseria* (e.g., type IV pili) suggest that they function as opportunistic pathobionts, or commensals capable of contributing to disease under permissive environmental conditions (12, 38–41). But despite their emerging clinical relevance, the physiological requirements that enable *Neisseria* to colonize the lower airway remain poorly defined.

A central knowledge gap is whether oral *Neisseria* employ distinct metabolic strategies to persist within the altered nutritional landscape of the inflamed lung. In CF and non-CF bronchiectasis, the lower airways are replete with host-derived substrates such as mucins, extracellular DNA, lactate, and nitrate, many of which accumulate as a consequence chronic inflammation and immune activation (42–47). While multi-omics analyses have revealed community-wide metabolic shifts involving these metabolites, few studies have examined how individual taxa, particularly commensals, exploit these resources to support their fitness in situ (12, 48, 49). Moreover, given that inflammation can remodel the airway environment in predictable ways, understanding how oral commensals respond to these changes could uncover broadly relevant principles of microbial persistence and inform new therapeutic approaches beyond conventional antibiotics, which are increasingly limited due to rising resistance and off-target disruption of beneficial microbiota (50).

Here, we used *N. mucosa*, one of the most abundant and widespread species of oral *Neisseria* (14, 36), as a model organism to dissect the metabolic requirements for oral commensal survival in the lower airway. To this end, we leveraged synthetic cystic fibrosis medium (SCFM2), a highly validated, chemically defined medium that mimics the nutrient composition of CF airway secretions (51–55). Using this in vitro system, we performed transposon sequencing (Tn-seq) (56) to identify genes required for *N. mucosa* fitness under anoxic, sputum-like conditions. This genome-wide screen revealed several critical metabolic functions, including L-lactate catabolism, pyrimidine biosynthesis, and nitrate respiration. Notably, while nitrate respiration is likely adaptive in inflamed contexts (57), including bronchiectasis (46, 58, 59), we show that it also supports *N. mucosa* growth in saliva, suggesting that commensal metabolic traits may be co-opted during opportunistic infection. Finally, we demonstrate that *N. mucosa* exhibits enhanced resistance to multiple, clinically relevant antibiotics when cultured in SCFM2, but remains susceptible to a selective inhibitor of nitrate respiration (60), highlighting the value of using disease-relevant media in therapeutic screening.

Together, our findings illuminate the metabolic logic by which a normally benign oral commensal can persist in the inflamed airway and underscore the therapeutic potential of targeting environment-specific microbial vulnerabilities in chronic lung disease.

## Results

### Synthetic sputum enhances *N. mucosa* antibiotic resistance and anoxic growth

*Neisseria* species are common colonists of the upper respiratory tract, including the oral cavity, in healthy individuals. However, to persist in the lower respiratory tract, particularly in patients with chronic airway diseases, commensal *Neisseria* must withstand numerous host and environmental stressors, including antibiotic exposure. The macrolide azithromycin is one of the most widely prescribed antibiotics for bronchiectasis (61, 62). It was also once a frontline treatment for pathogenic *Neisseria* (e.g., *N. gonorrhoeae*), but is now largely avoided due to rising resistance, which can emerge in both pathogenic and commensal species (63, 64). While resistance is often linked to genetic mechanisms (e.g., target site mutation), non-heritable “phenotypic” antibiotic resistance, driven by environmental factors such as pH or nutrient availability, can also contribute to reduced susceptibility (65, 66).

To explore this in *N. mucosa*, a common commensal and opportunistic lung colonist, we used synthetic cystic fibrosis medium (SCFM2), which mimics the nutrient composition of sputum from individuals with cystic fibrosis, a condition often associated with bronchiectasis (51). Using a strain originally isolated from patient sputum (ATCC 19696 (67)), we determined the minimum inhibitory concentration (MIC) of azithromycin on SCFM2 and compared it to that on a standard nutrient-rich laboratory medium, tryptic soy broth supplemented with yeast extract (TSBYE). On TSBYE, the MIC of azithromycin was 4 µg/mL, consistent with reported values for resistant clinical isolates (68) (**Fig. 1A, B**). In contrast, the MIC on SCFM2 exceeded 256 µg/mL, indicating dramatically reduced susceptibility under sputum-like conditions (**Fig. 1A, B**).

**Figure 1.**
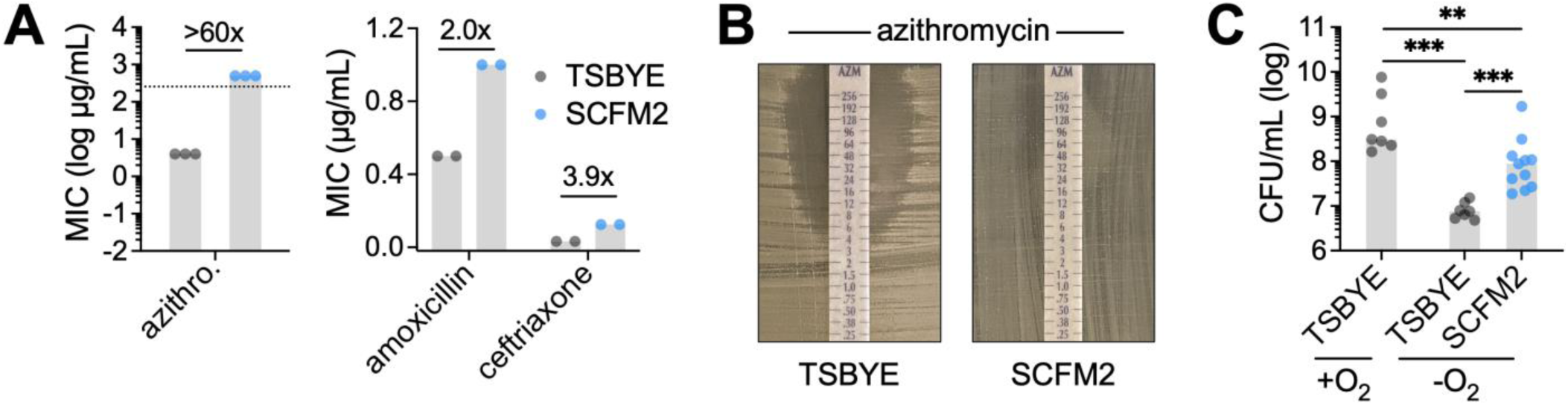
Synthetic sputum enhances *N. mucosa* antibiotic resistance and anoxic growth. (**A**) Minimum inhibitory concentrations (MICs) of the indicated antibiotics for *N. mucosa* cultured as a lawn on tryptic soy broth + yeast extract (TSBYE) or synthetic cystic fibrosis medium (SCFM2) agar. Lawns were cultured under oxic conditions since growth was not perceptible under anoxic conditions. Data represent ≥2 biological replicates (performed on separate days); gray bars, median; numbers above bars, fold change in median. Dotted line (left panel) indicates the max MIC value. (**B**) Representative MIC results for azithromycin (AZM). (**C**) Growth yields of *N. mucosa* after culture for 24 h in TSBYE or SCFM2 liquid media under oxic (+O_2_) or anoxic (-O_2_) conditions. Media were not de-oxygenated prior to inoculation at OD_600_ = 0.001 (∼10^5^ CFU/mL). Data represent ≥4 biological replicates, each with 1-2 technical replicates (total n = 7-11); gray bars, median; **, P < 0.01; ***, P < 0.001 (two-tailed Mann-Whitney test).

To extend this analysis, we determined the MICs of two additional, clinically relevant antibiotics: the β-lactam amoxicillin, commonly used to treat bronchiectasis (69), and the cephalosporin ceftriaxone, a standard treatment for invasive *Neisseria* infections (63). In both cases, *N. mucosa* exhibited moderately increased resistance on SCFM2 compared to TSBYE (**Fig. 1A**), suggesting a broader pattern of phenotypic resistance in sputum-like environments.

Given that sputum is often anoxic (70, 71), we next examined the impact of oxygen limitation on *N. mucosa* growth. As expected based on the frequent classification of *Neisseria* as “aerobes” (72), *N. mucosa* exhibited a >40-fold lower growth yield under anoxic versus oxic conditions in TSBYE (**Fig. 1C**). In contrast, when cultured under the same anoxic conditions in SCFM2, *N. mucosa* achieved a final yield 12-fold higher than in TSBYE (**Fig. 1C**), suggesting improved adaptation to oxygen limitation.

Together, these findings indicate that sputum-like conditions modeled by SCFM2 promote both phenotypic antibiotic resistance and anoxic growth in *N. mucosa*. These results highlight how local environmental factors at the infection site can potentially enhance the fitness and persistence of commensal *Neisseria* in the lower respiratory tract.

### Tn-seq identifies *N. mucosa* fitness determinants in synthetic sputum

Given that SCFM2 promotes phenotypic antibiotic resistance in *N. mucosa*, we next sought to identify bacterial functions that are essential for growth in this sputum-like environment. In doing so, we aimed to uncover potential targets for novel, non-antibiotic therapies that could directly disrupt *N. mucosa* fitness in the lung.

To this end, we performed a genome-wide transposon (Tn) mutant screen, Tn-seq, to identify genes required for *N. mucosa* replication in SCFM2. A key advantage of Tn-seq is that it enables rapid quantification of relative Tn mutant abundance (fitness) across a pooled Tn mutant library by sequencing genomic regions adjacent to Tn insertions (56).

To generate a Tn mutant pool in *N. mucosa*, we leveraged its natural competence for transformation (73). Specifically, we mutagenized purified *N. mucosa* genomic DNA in vitro using EZ-Tn5 transposition (74) and then naturally transformed this DNA back into *N. mucosa* cells (see Methods). The resulting transformants were pooled and preserved as single-use aliquots. Analysis of three independent aliquots identified 33,568 unique high-confidence Tn insertions (i.e., present in all three replicates), corresponding to an average of one insertion every 80 bp across the ∼2.7 Mbp *N. mucosa* 19696 genome.

We next subjected the Tn mutant pool to anoxic growth in SCFM2 and performed Tn-seq (**Fig. 2A**). Comparing mutant abundance before and after growth (input vs. SCFM2 condition) revealed 46 genes with significantly reduced representation (P_adj_ < 0.1), suggesting that these genes are required for *N. mucosa* fitness in SCFM2 (**Fig. 2B**, left side of plot). Among these, three genes exhibited particularly strong depletion: 1) an oxidoreductase of unknown function, 2) *tamB*, encoding a subunit of the translocation and assembly module (TAM) complex involved in outer membrane protein insertion (75), and 3) *mrdA*, encoding a peptidoglycan transpeptidase essential for cell wall synthesis (76).

**Figure 2.**
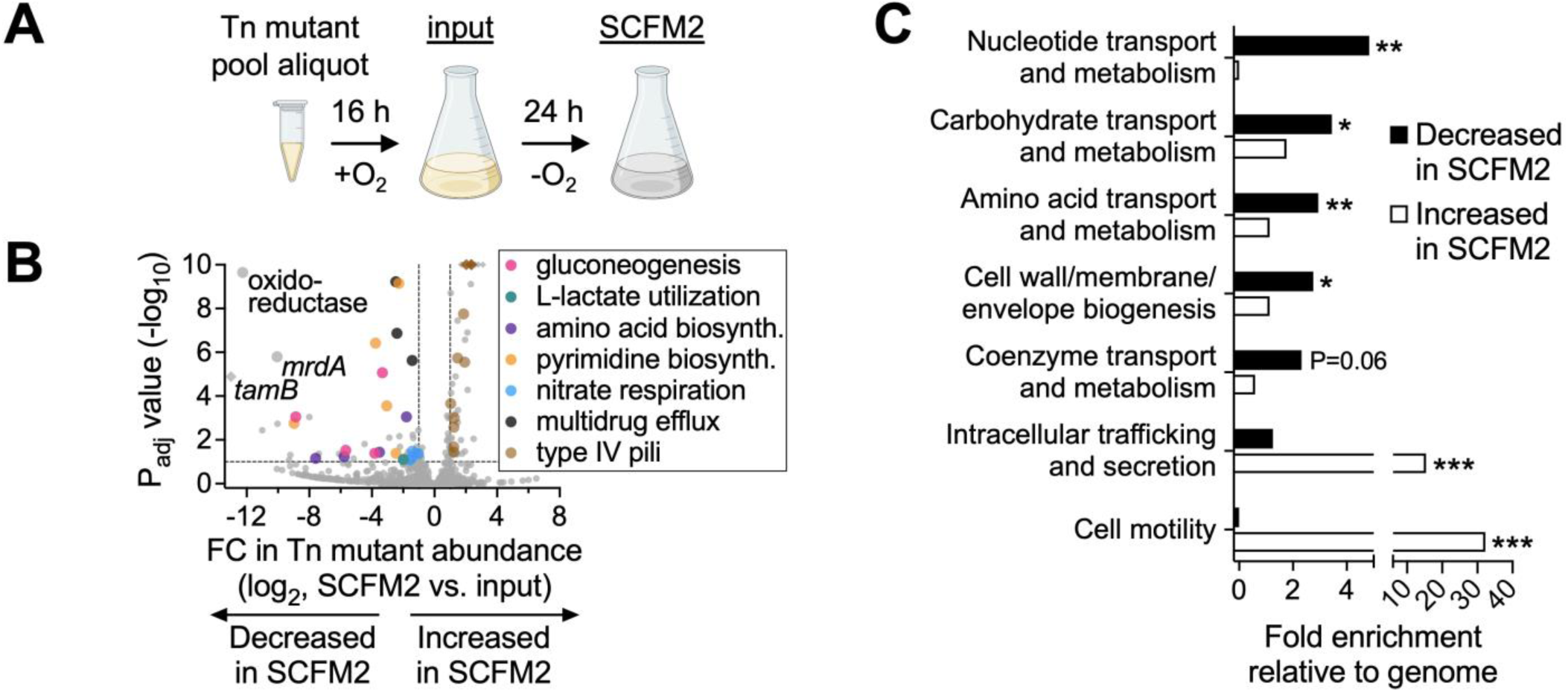
Tn-seq identifies *N. mucosa* fitness determinants in synthetic sputum. (**A**) Transposon (Tn)-seq experimental design. A cryo-aliquot of the *N. mucosa* Tn mutant pool was revived for 16-20 h in tryptic soy broth + yeast extract (TSBYE) under oxic conditions (input condition), then back-diluted and cultured for 24 h in synthetic cystic fibrosis medium under anoxic conditions (SCFM2 condition). Media were not de-oxygenated prior to inoculation. (**B**) Differential abundance of *N. mucosa* Tn mutants after anoxic growth in SCFM2. Genes (points) are colored by functional category. The x-axis represents fold change (FC) in total Tn mutant abundance per gene (SCFM2 vs. input); y-axis, adjusted P values (DESeq2; Wald test with Benjamini-Hochberg correction); dashed lines, significance cutoffs (FC > 2; P_adj_ < 0.1 [default DESeq2 setting]). For visual clarity, 13 off-axis values (diamonds) were adjusted to the plot’s maximum FC or P value. (**C**) Enrichment of clusters of orthologous groups (COG) categories among genes with decreased or increased Tn mutant abundance following anoxic growth in SCFM2. The x-axis represents the fraction of SCFM2-decreased or -increased genes in a COG category divided by the genome-wide fraction of genes in the same category. *, P < 0.05; **, P < 0.01; ***, P < 0.001 (one-sided hypergeometric test). Only enriched COG categories with P < 0.07 are shown.

To identify functional trends within this gene set, we performed an enrichment analysis using clusters of orthologous groups (COGs) (77). Of the 21 COG categories analyzed, four were significantly overrepresented among the SCFM2-depleted genes relative to their distribution across the genome (P < 0.05) (**Fig. 2C**). These included “Cell wall/membrane/envelope biogenesis,” which encompasses *mrdA*, and three metabolic categories: “Carbohydrate transport and metabolism,” “Amino acid transport and metabolism,” and “Nucleotide transport and metabolism.”

Together, these findings indicate that metabolic flexibility and maintenance of envelope integrity are critical for *N. mucosa* replication under sputum-like conditions. These processes may represent key vulnerabilities for therapeutic exploitation in the context of chronic airway colonization.

### Gluconeogenesis and L-lactate catabolism promote *N. mucosa* fitness in synthetic sputum

The enrichment of the COG category “Carbohydrate transport and metabolism” was driven by four genes involved in glucose metabolism: *fba* (fructose 1,6-bisphosphate aldolase), *pgk* (phosphoglycerate kinase), *gpmA* (phosphoglycerate mutase), and *ppsA* (phosphoenolpyruvate synthase) (**Fig. 2B**, pink points). As in other *Neisseria* species, *N. mucosa* likely catabolizes glucose via the Entner-Doudoroff (ED) pathway, as it lacks phosphofructokinase (Pfk), a key enzyme in the Embden-Meyerhof-Parnas (EMP) pathway (40) (**Fig. 3A**). In the ED pathway, glucose is first converted to pyruvate and glyceraldehyde 3-phosphate (Ga3P), which is then further metabolized into pyruvate via enzymes shared with the lower part of the EMP pathway (78) (**Fig. 3A**). Under glucose-limited conditions, much of the EMP pathway, including Fba, Pgk, and GpmA, can operate in reverse to support gluconeogenesis, regenerating Ga3P and upstream intermediates for anabolic processes.

**Figure 3.**
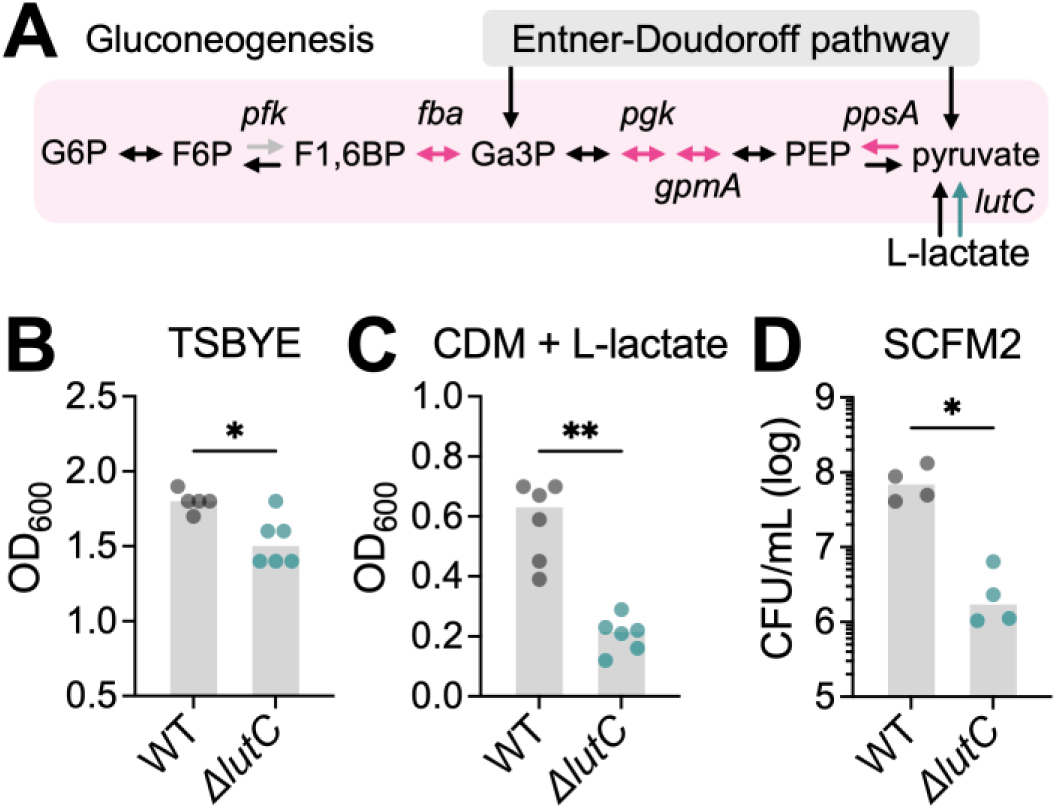
Gluconeogenesis and L-lactate catabolism promote *N. mucosa* fitness in synthetic sputum. (**A**) *N. mucosa* carbon utilization pathways. Enzymes (arrows) highlighted in pink and teal were identified by Tn-seq as fitness determinants in synthetic cystic fibrosis medium (SCFM2). Genes: *pfk*, phosphofructokinase (not present in *N. mucosa* genome); *fba*, fructose 1,6-bisphosphate (F1,6BP) aldolase; *pgk*, phosphoglycerate kinase; *gpmA*, phosphoglycerate mutase; *ppsA*, phosphoenolpyruvate (PEP) synthase; *lutC*, L-lactate dehydrogenase. Metabolites: G6P, glucose 6-phosphate; F6P, fructose 6-phosphate; Ga3P, glyceraldehyde 3-phosphate. (**B-D**) Growth yields of the *N. mucosa* wild type (WT) and L-lactate dehydrogenase mutant (*ΔlutC*) after culture for 24 h in (B) tryptic soy broth + yeast extract (TSBYE) under oxic conditions, (C) Socransky’s chemically defined medium (CDM) + 20 mM sodium L-lactate under oxic conditions, or (D) SCFM2 under anoxic conditions. Media were not de-oxygenated prior to inoculation at OD_600_ = 0.001 (∼10^5^ CFU/mL). CFU were necessary to assess growth in SCFM2 due to its turbidity. Data represent ≥2 biological replicates (performed on separate days), each with ≥2 technical replicates (total n = 4-6); gray bars, median; *, P < 0.05; **, P < 0.01 (two-tailed Mann-Whitney test).

In many respiratory pathogens, glucose can serve as a key carbon source in artificial sputum (79, 80). In contrast, despite the presence of 3 mM glucose in SCFM2, *N. mucosa* appears to rely more heavily on gluconeogenesis than the ED pathway for fitness in this environment. Supporting this, none of the enzymes specific to the ED pathway were identified as fitness determinants. However, both phosphoenolpyruvate synthase (PpsA), a strictly gluconeogenic enzyme, and fructose-1,6-bisphosphate aldolase (Fba), a reversible enzyme upstream of Ga3P that does not overlap with the ED pathway, were essential for *N. mucosa* fitness in SCFM2 (**Fig. 3A**).

L-lactate is a gluconeogenic substrate and key in vivo carbon source for pathogenic *Neisseria* (81, 82). Like its pathogenic relatives, *N. mucosa* encodes at least two membrane-bound, respiratory L-lactate dehydrogenases: LldD and LutACB (83). Both enzymes oxidize L-lactate to pyruvate while coupling this reaction to cellular respiration, contributing to ATP synthesis via oxidative phosphorylation. Despite the apparent redundancy of these enzymes, only LutACB (specifically LutC) was identified by Tn-seq as a fitness determinant in SCFM2 (**Fig. 2B**, teal point).

To validate these findings, we constructed a *lutC* deletion mutant. In nutrient-rich TSBYE, the mutant displayed only a mild growth defect (1.2-fold reduction in final yield) compared to the wild type (WT) (**Fig. 3B**). As expected, this defect was more pronounced (2.9-fold reduction) in a chemically defined medium (CDM) with L-lactate as the sole carbon source (**Fig. 3C**). Most strikingly, the *ΔlutC* mutant exhibited a 40-fold reduction in yield relative to the WT in SCFM2 (**Fig. 3D**), providing strong evidence that L-lactate is a major carbon source for *N. mucosa* under sputum-like conditions.

Given that L-lactate is a gluconeogenic substrate, these findings support the following model for *N. mucosa* carbon utilization in SCFM2: (1) the abundant L-lactate present in this environment (9 mM) is oxidized to pyruvate via LutACB; (2) this pyruvate is then utilized via gluconeogenesis to generate biosynthetic precursors and support growth (**Fig. 3A**).

### Amino acid and pyrimidine biosynthesis support *N. mucosa* fitness in synthetic sputum

We next examined the COG category “Amino acid transport and metabolism” (**Fig. 2C**), which was enriched primarily due to six genes involved in the biosynthesis of distinct amino acids or amino acid families: *dapF* (lysine), *proC* (proline), *thrC* (threonine), *aspA* (aspartate), *hisG* (histidine), and *ilvB* (branched-chain amino acids [BCAA] isoleucine, leucine, and valine) (**Fig. 2B**, purple points).

Surprisingly, all eight amino acids associated with these genes are present in SCFM2, with six of the eight exceeding the median amino acid concentration (**Fig. 4A**). This suggests that, despite their presence, amino acids levels in SCFM2 are not sufficient to meet *N. mucosa*’s metabolic demands, necessitating de novo synthesis. Notably, *dapF* contributes not only to lysine biosynthesis but also to the production of meso-2,6-diaminopimelate, a key precursor for peptidoglycan (76). Its essentiality may therefore reflect dual roles in supporting both amino acid and cell wall biosynthesis, consistent with the enrichment of the “Cell wall/membrane/envelope biogenesis” COG category (**Fig. 2C**).

**Figure 4.**
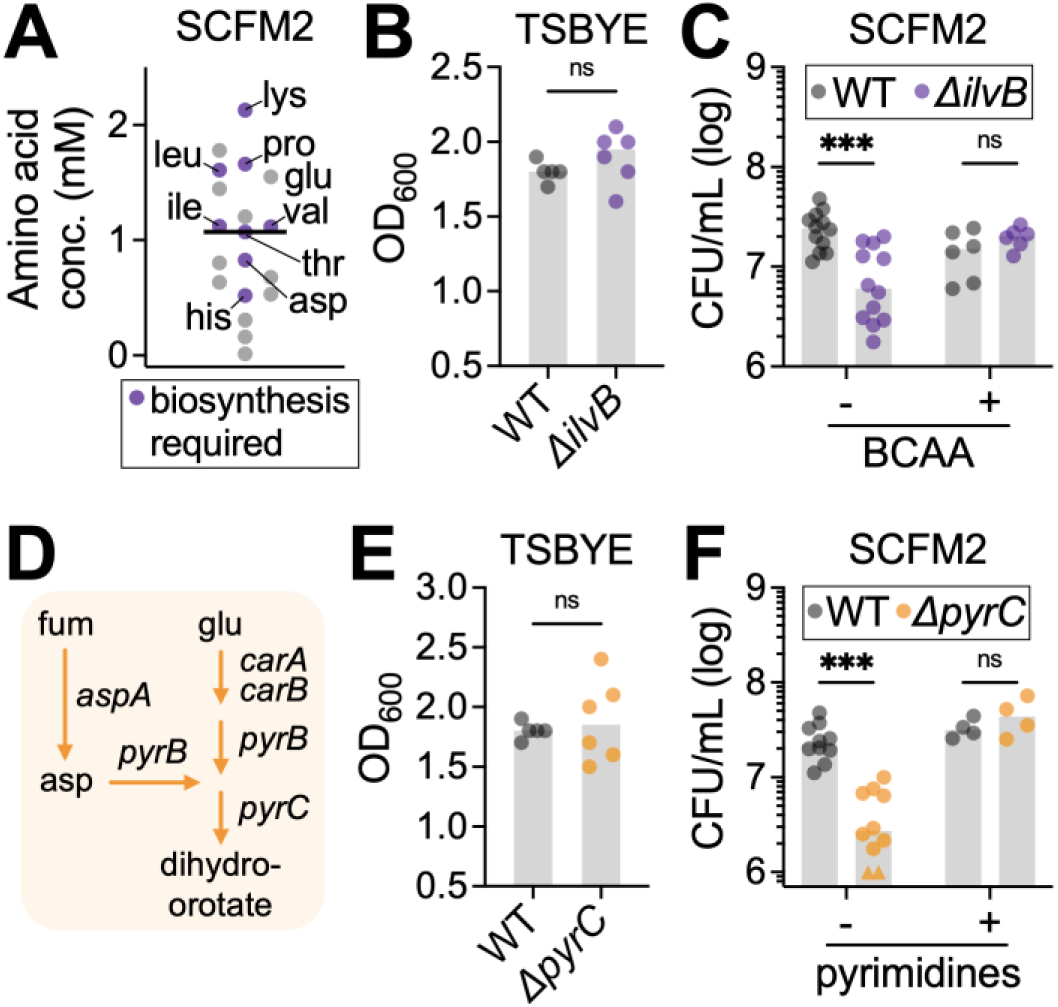
Amino acid and pyrimidine biosynthesis support *N. mucosa* fitness in synthetic sputum. (**A**) Concentration of amino acids in synthetic cystic fibrosis medium (SCFM2). Purple indicates at least one biosynthetic gene was identified by Tn-seq as required for *N. mucosa* fitness in SCFM2. Abbreviations: lys, lysine; pro, proline; glu, glutamate; ile, isoleucine; leu, leucine; val, valine; thr, threonine; asp, aspartate; his, histidine. (**B, C, E, F**) Growth yields of the *N. mucosa* wild type (WT), acetolactate synthase mutant (*ΔilvB*), and dihydroorotase mutant (*ΔpyrC*) after culture for 24 h in (B, E) tryptic soy broth + yeast extract (TSBYE) under oxic conditions or (C, F) SCFM2 under anoxic conditions. Where indicated, SCFM2 was supplemented with the branched-chain amino acids (BCAA) isoleucine, leucine, and valine (each 10 mM), the pyrimidines cytosine, thymine, and uracil (each 0.2 mM), or an equal volume of H_2_O (vehicle). Media were de-oxygenated in (C) and (F), but not (B) or (E), prior to inoculation at OD_600_ = 0.001 (∼10^5^ CFU/mL). CFU were necessary to assess growth in SCFM2 due to its turbidity. Data represent ≥2 biological replicates (performed on separate days), each with ≥2 technical replicates (total n = 4-12); gray bars, median; ***, P < 0.001; ns, not significant (two-tailed Mann-Whitney test). (F) For visual clarity, 2 below-axis values (triangles) were adjusted to 10^6^ CFU/mL. (**D**) *N. mucosa* biosynthetic pathways for the pyrimidine precursor dihydroorotate. All depicted enzymes (arrows) were identified by Tn-seq as fitness determinants in SCFM2. Genes: *carA* and *B*, carbamoyl phosphate synthase small and large subunits; *pyrB*, aspartate carbamoyltransferase; *pyrC*, dihydroorotase; *aspA*, aspartate ammonia-lyase. Metabolites: fum, fumarate; asp, aspartate; glu, glutamate.

To validate the role of amino acid biosynthesis, we constructed a deletion mutant of *ilvB*, which encodes a subunit of acetolactate synthase, the first committed step in BCAA biosynthesis (84). While this mutant grew comparably to the WT in nutrient-rich TSBYE (**Fig. 4B**), it exhibited a 4-fold reduction in yield in SCFM2, consistent with Tn-seq results (**Fig. 4C**). Supplementing SCFM2 with exogenous BCAA fully restored the mutant’s growth to WT levels, confirming that limited BCAA availability constrains *N. mucosa* fitness in this environment (**Fig. 4C**).

The parallel enrichment of the COG category “Nucleotide transport and metabolism” (**Fig. 2C**) was driven primarily by four genes involved in pyrimidine biosynthesis: *carA*, *carB*, *pyrB*, and *pyrC* (**Fig. 2B**, orange points). This pathway uses L-glutamate and L-aspartate, both present in SCFM2 (**Fig. 4A**, labeled “glu” and “asp”), to synthesize dihydroorotate, a central intermediate in pyrimidine biosynthesis (85) (**Fig. 4D**). In addition, *aspA*, which synthesizes L-aspartate from fumarate, was also required, linking pyrimidine biosynthesis to central carbon metabolism (**Fig. 4D**).

Although SCFM2 contains DNA as a structural component, it lacks free pyrimidines (cytosine, thymine, and uracil), likely explaining *N. mucosa*’s strong reliance on de novo synthesis. To validate this, we deleted *pyrC*, which encodes dihydroorotase (**Fig. 4D**). As with the *ΔilvB* mutant, the *ΔpyrC* mutant grew normally in TSBYE (**Fig. 4E**), but exhibited an 8-fold reduction in SCFM2 (**Fig. 4F**). This defect was fully rescued by exogenous pyrimidines, confirming that pyrimidine limitation restricts *N. mucosa* growth under sputum-like conditions (**Fig. 4F**).

Together, these findings highlight the nutritional constraints that are imposed on *N. mucosa* in SCFM2. Despite the presence of amino acids and nucleotides, their concentrations—or bioavailable forms—are insufficient to support optimal growth, driving *N. mucosa* to perform de novo synthesis to meet its metabolic needs.

### Nitrate respiration promotes *N. mucosa* fitness in synthetic sputum

The near-significant enrichment of the COG category “Coenzyme transport and metabolism” (P = 0.06) was primarily driven by genes involved in the biosynthesis of thiamine (*thiE*), glutathione (*gshA*), and molybdopterin (*mogA*) (**Fig. 2C**). Of these, molybdopterin was of particular interest, as it is an essential cofactor for anaerobic reductases (60) (**Fig. 5A**). These enzymes enable respiration using alternative terminal electron acceptors under anoxic conditions, suggesting that anaerobic respiration may contribute to *N. mucosa*’s enhanced growth in SCFM2 under oxygen-limited conditions (**Fig. 1C**). Supporting this, *N. mucosa* encodes genes for denitrification, a respiratory pathway that reduces nitrate (NO_3_ ^-^) to nitrogen gas (N_2_) (86), and two such genes, *narG* (nitrate reductase) and *norB* (nitric oxide reductase), were identified as fitness determinants in SCFM2 (**Fig. 5A**).

**Figure 5.**
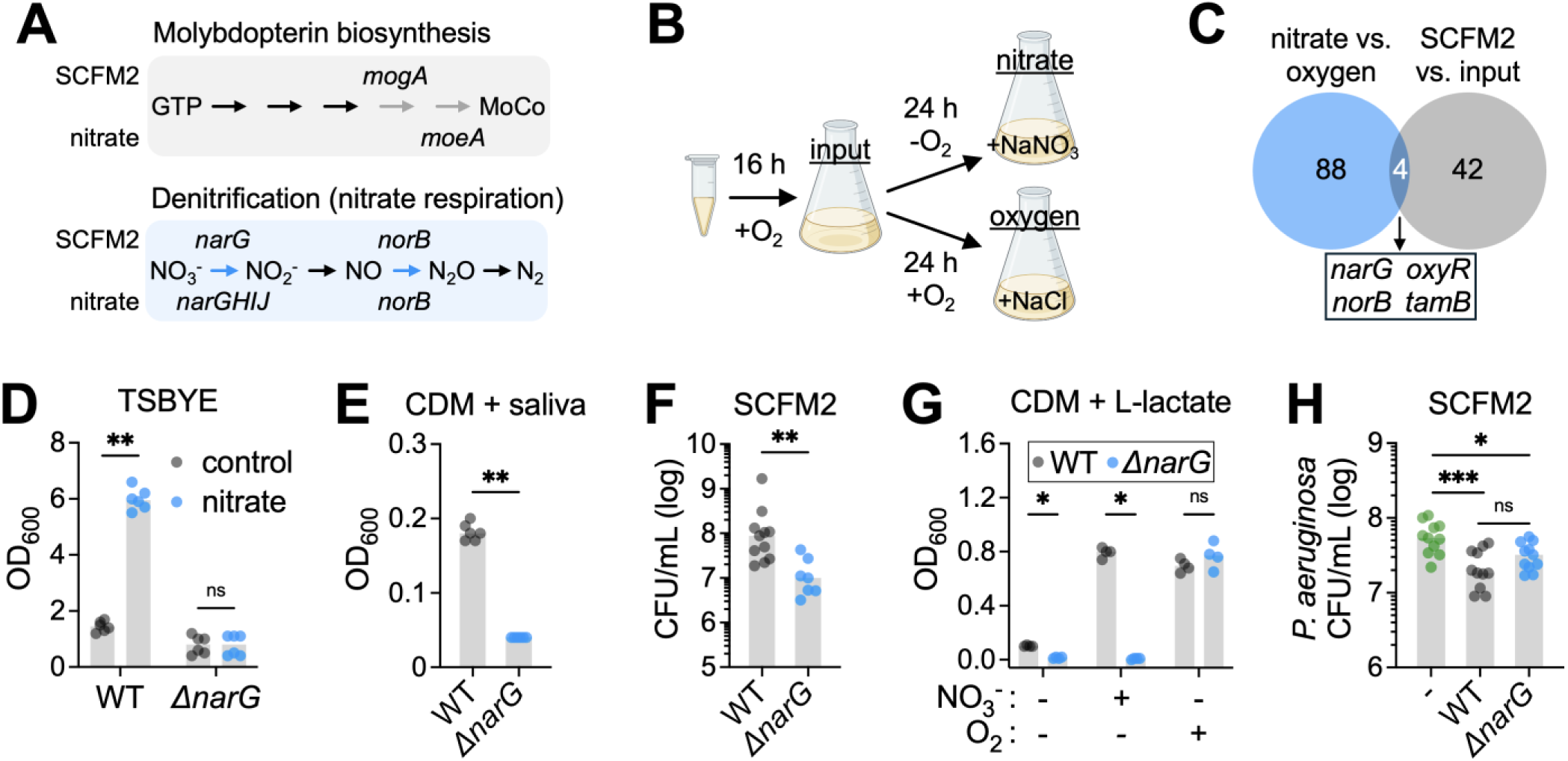
Nitrate respiration promotes *N. mucosa* fitness in synthetic sputum. (**A**) *N. mucosa* molybdopterin biosynthesis and denitrification pathways. Enzymes (arrows) highlighted in gray and blue were identified by Tn-seq as fitness determinants in synthetic cystic fibrosis medium (SCFM2) or nitrate-supplemented tryptic soy broth + yeast extract. Gene names above arrows: significant in SCFM2. Gene names below arrows: significant in nitrate-supplemented TSBYE. Genes: *mogA*, molybdopterin adenylyltransferase; *moeA*, molybdopterin molybdenumtransferase; *narG*, nitrate reductase alpha (catalytic) subunit; *narHIJ*, nitrate reductase beta subunit, gamma subunit, and chaperone protein; *norB*, nitric oxide reductase catalytic subunit. Metabolites: GTP, guanosine triphosphate; MoCo, molybdenum cofactor; NO_3_^-^, nitrate; NO_2_^-^, nitrite; NO, nitric oxide; N_2_O, nitrous oxide; N_2_, nitrogen gas. Tn-seq results for *moeA*, *narH*, *narI*, and *narJ* were marginally below the fold-change cutoff for significance (P_adj_ < 0.03, but FC only 1.8-2). (**B**) Transposon (Tn)-seq experimental design. A cryo-aliquot of the *N. mucosa* Tn mutant pool was revived for 16-20 h in TSBYE under oxic conditions (input condition), then back-diluted and cultured for 24 h in TSBYE + 10 mM NaNO_3_ under anoxic conditions (nitrate condition) or TSBYE + 10 mM NaCl under oxic conditions (oxygen condition). Media were de-oxygenated prior to inoculation. (**C**) Venn diagram showing the overlap between the number of *N. mucosa* fitness determinants in TSBYE + nitrate (when compared to TSBYE + oxygen) and SCFM2 (when compared to the input). (**D-G**) Growth yields of the *N. mucosa* wild type (WT) and nitrate reductase mutant (*ΔnarG*) after culture for 24 h in (D) TSBYE + 10 mM NaCl (control) or NaNO_3_ (nitrate) under anoxic conditions, (E) Socransky’s chemically defined medium (CDM) + 25% saliva (pooled from healthy human donors) under anoxic conditions, (F) SCFM2 under anoxic conditions, or (G) CDM + 20 mM sodium L-lactate under the indicated conditions (-O_2_, anoxic; +O_2_, oxic; -NO_3_^-^, +10 mM NaCl; +NO_3_^-^, +10 mM NaNO_3_). Media were not de-oxygenated prior to inoculation at OD_600_ = 0.001 (∼10^5^ CFU/mL). CFU were necessary to assess growth in SCFM2 due to its turbidity. (F) The WT data is the same as the SCFM2 data in Fig. 1C. (**H**) Growth yields of *P. aeruginosa* PA14 after culture for 24 h in SCFM2 under anoxic conditions in mono- or co-culture with the *N. mucosa* WT or *ΔnarG* mutant. Media were de-oxygenated prior to inoculation with ∼10^3^ CFU/mL for *P. aeruginosa* and ∼10^5^ CFU/mL for *N. mucosa*. (D-H) Data represent ≥2 biological replicates (performed on separate days), each with 1-4 technical replicates (total n = 4-11); gray bars, median; *, P < 0.05; **, P < 0.01; ***, P < 0.001; ns, not significant (two-tailed Mann-Whitney test).

To more directly assess genes required for growth with nitrate, we performed additional Tn-seq experiments under defined conditions. Specifically, we compared growth in nutrient-rich TSBYE under anoxic conditions with nitrate as the terminal electron acceptor versus growth under oxic conditions with oxygen as the acceptor (**Fig. 5B**). This analysis identified 92 genes that are specifically required for nitrate-supported anaerobic growth but dispensable in the presence of oxygen (**Fig. 5C**).

Of these 92 genes, four overlapped with SCFM2 fitness determinants, representing a trend toward significant enrichment (P = 0.056, one-sided hypergeometric test) (**Fig. 5C**). As expected, two of these genes, *narG* and *norB*, encode core components of the denitrification pathway (**Fig. 5A**). A third, *oxyR*, encodes a transcriptional regulator involved in oxidative and nitrosative stress responses, potentially related to nitric oxide-induced toxicity during denitrification (87). The fourth gene, *tamB*, encodes a subunit of the translocation and assembly module (TAM) complex required for outer membrane protein assembly (75). Notably, *tamB* exhibited the most pronounced depletion among all SCFM2 fitness determinants (log_2_ fold change < -22; **Fig. 2B**), further linking nitrate respiration to broader fitness requirements under sputum-like conditions.

To validate the role of nitrate respiration, we constructed a *narG* deletion mutant. Unlike the WT, the *ΔnarG* mutant failed to achieve enhanced growth in TSBYE supplemented with nitrate (**Fig. 5D**). The mutant was also attenuated in a chemically defined medium (CDM) supplemented with pooled human saliva, a physiologically relevant nutrient source in the oral cavity, where nitrate can temporarily reach levels as high as 5-10 mM following nitrate-rich meals (88) (**Fig. 5E**). Most notably, the *ΔnarG* mutant exhibited a 9-fold reduction in yield compared to the WT in SCFM2, consistent with Tn-seq results (**Fig. 5F**).

Given that L-lactate is a key non-fermentable carbon source for *N. mucosa* in SCFM2 (**Fig. 3**), we next tested whether nitrate respiration enables L-lactate utilization under anoxic conditions. Indeed, the *ΔnarG* mutant showed a 7-fold reduction in anoxic growth yield in CDM supplemented with L-lactate, which increased to an 80-fold defect with added nitrate (10 mM; **Fig. 5G**). These defects were specific to anaerobic metabolism, as the mutant grew comparably to the WT under oxic conditions (**Fig. 5G**).

In the inflamed airway, *N. mucosa* likely faces competition from other nitrate-respiring bacteria, particularly *Pseudomonas aeruginosa*, which also depends on nitrate respiration for fitness in synthetic sputum (58). To assess the impact of nitrate respiration on interspecies competition, we compared *P. aeruginosa* growth in mono- and co-culture with either the *N. mucosa* WT or *ΔnarG* mutant in SCFM2 (**Fig. 5H**). Co-culture with the WT significantly reduced *P. aeruginosa* yield (2.9-fold), whereas co-culture with the *ΔnarG* mutant had a more modest effect (1.7-fold), suggesting that nitrate respiration contributes to *N. mucosa*’s competitive fitness in polymicrobial settings (**Fig. 5H**).

Together, these results demonstrate that nitrate respiration is a key fitness determinant for *N. mucosa* under sputum-like conditions. It not only supports anoxic growth on L-lactate but also enhances competitive interactions with other nitrate-utilizing airway pathogens.

### Targeting *N. mucosa* metabolic vulnerabilities in synthetic sputum

Having identified key genes required for *N. mucosa* fitness in SCFM2, we next explored whether any of these metabolic functions could be selectively disrupted using non-antibiotic compounds. A major motivation was our observation that *N. mucosa* displays reduced susceptibility to antibiotics under sputum-like conditions (**Fig. 1A, B**), potentially contributing to the limited efficacy of conventional antibiotic therapies.

We first examined *N. mucosa*’s dependence on L-lactate catabolism via the L-lactate dehydrogenase complex LutACB, which was a strong fitness determinant in SCFM2 (**Fig. 3D**). Treatment with oxalate, a known inhibitor of L-lactate dehydrogenases (89), impaired *N. mucosa* growth in SCFM2 but not in nutrient-rich TSBYE (**Fig. S1A, B**), consistent with the differential requirement for *lutC* in these environments (**Fig. 3B, D**). However, in mixed-strain competition experiments, oxalate treatment failed to selectively inhibit the WT, which still robustly outcompeted the *ΔlutC* mutant even in the presence of oxalate (**Fig. S1C**). This result suggests that oxalate exerts broader, non-selective inhibitory effects in SCFM2, reducing its utility for targeting L-lactate metabolism.

We next tested whether inhibition of de novo pyrimidine biosynthesis could selectively impair *N. mucosa* in SCFM2. We focused on the small molecule PALA (N-phosphonacetyl-L-aspartate), an inhibitor of aspartate carbamoyltransferase (*pyrB*), which was required for fitness in SCFM2 (**Fig. 4D**). Consistent with PALA’s reported context-dependent activity (90), PALA inhibited *N. mucosa* growth in SCFM2 but not TSBYE (**Fig. S1D, E**). However, as with oxalate, PALA failed to selectively inhibit the WT. In competition assays, the *ΔpyrC* mutant (which should be resistant to PALA due to its downstream position in the pathway) was outcompeted by the WT even under PALA treatment (**Fig. S1F**), suggesting PALA may also have off-target or general inhibitory effects in this context.

Lastly, we investigated whether nitrate respiration, another essential function in SCFM2 (**Fig. 5F**), could be disrupted using tungstate (WO_4_^2-^), an analog of molybdate (MO_4_^2-^) that inhibits molybdo-enzymes such as nitrate reductase (60). In TSBYE, tungstate at 100 µM impaired growth under nitrate-respiring conditions while enhancing growth under aerobic conditions, indicating selective inhibition of nitrate reductase (**Fig. 6A, B**). At higher concentrations (1 mM), tungstate impaired growth under both conditions, likely due to broader metabolic disruption (**Fig. 6A**). Based on this, 100 µM tungstate was used for subsequent assays. In SCFM2, this concentration strongly suppressed *N. mucosa* growth under anoxic conditions (**Fig. 6C**), and critically, abolished the competitive advantage of the WT over the *ΔnarG* mutant (**Fig. 6D**). To determine whether this strategy might extend to other airway pathogens, we tested tungstate against *P. aeruginosa*. As observed with *N. mucosa*, *P. aeruginosa* growth in SCFM2 was significantly impaired by tungstate (**Fig. 6E**).

**Figure 6.**
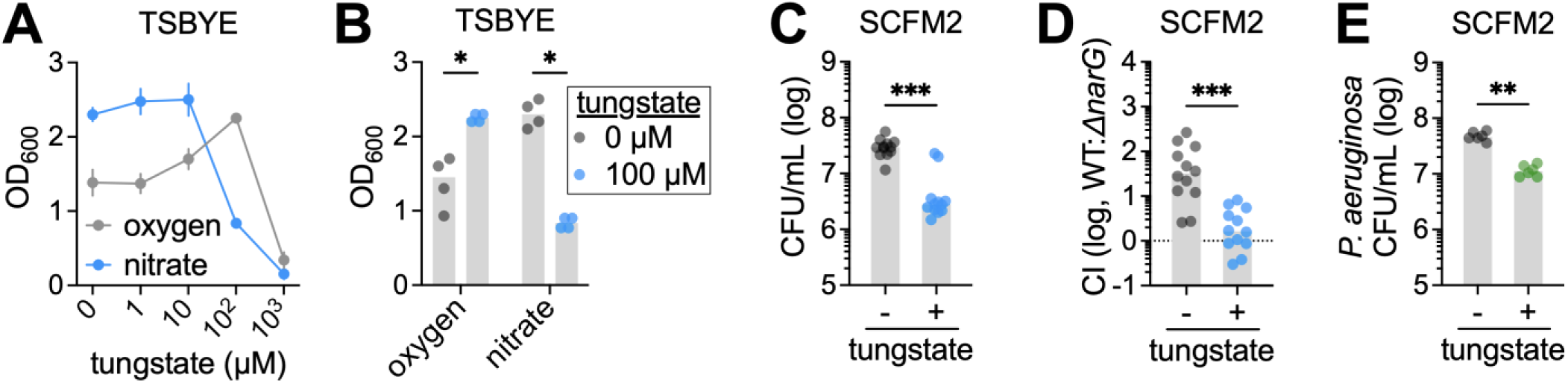
Tungstate selectively inhibits *N. mucosa* nitrate respiration in synthetic sputum. (**A, B**) Growth yields of *N. mucosa* after culture for 24 h in tryptic soy broth + yeast extract (TSBYE) with increasing amounts of tungstate. Oxygen: TSBYE + 10 mM NaCl incubated under oxic conditions. Nitrate: TSBYE + 10 mM NaNO_3_ incubated under anoxic conditions. For visual clarity, statistical comparisons are only shown in (B) for 0 and 100 µM sodium tungstate. (**C, E**) Growth yields of (C) *N. mucosa* and (E) *P. aeruginosa* PA14 after culture for 24 h under anoxic conditions in synthetic cystic fibrosis medium (SCFM2) + 100 µM sodium tungstate or an equal volume of H_2_O (-). (**D**) Competitive indexes (CI) for the *N. mucosa* wild type (WT) and nitrate reductase mutant (*ΔnarG*) after co-culture for 24 h under anoxic conditions in SCFM2 + 100 µM sodium tungstate or an equal volume of H_2_O (-). CI determined by dividing the output WT:mutant ratio by the input WT:mutant ratio (∼1:1); a CI of 1 (log_10_CI = 0; dotted line) indicates equal fitness between WT and mutant. (A-E) Media were de-oxygenated in (C-E), but not (A) or (B), prior to inoculation at OD_600_ = 0.001 (∼10^5^ total CFU/mL). CFU were necessary to assess growth in SCFM2 due to its turbidity. Data represent ≥2 biological replicates (performed on separate days), each with ≥2 technical replicates (total n = 4-12); (A) points, mean ± standard error; (B-E) gray bars, median; *, P < 0.05; **, P < 0.01; ***, P < 0.001 (two-tailed Mann-Whitney test).

Together, these findings underscore the importance of studying microbial physiology in disease-relevant environments. Metabolic pathways that are dispensable in nutrient-rich media may be critical for survival in the inflamed airway, revealing context-specific vulnerabilities not apparent under standard laboratory conditions. Targeting such conditional dependencies offers a promising strategy to suppress opportunistic pathogens such as *N. mucosa*, particularly in settings where conventional antibiotics are less effective.

## Discussion

In this study, we identified key metabolic pathways that support the fitness of *N. mucosa* in synthetic cystic fibrosis medium (SCFM2), a validated in vitro model that mimics the nutrient environment of the chronically inflamed lower airway (51–55). Our results demonstrate that L-lactate catabolism, de novo amino acid and pyrimidine biosynthesis, and anaerobic nitrate respiration are essential for *N. mucosa* growth and survival under sputum-like conditions. These findings provide new mechanistic insight into how oral commensals metabolically adapt to disease-associated microenvironments (17–20).

*N. mucosa* and related commensal *Neisseria* species are typically regarded as health-associated taxa, predominating in the upper airway microbiota of healthy individuals, including the tongue and dental plaque (14, 36). However, growing evidence implicates these organisms in chronic lung diseases such as bronchiectasis, cystic fibrosis, and COPD (12, 13, 21–23, 37). Our data support this emerging view by suggesting that *N. mucosa* can engage metabolic programs, akin to those of pathogenic *Neisseria* (39, 40), to persist in the inflamed airway.

Specifically, we found that *N. mucosa* proliferates in SCFM2 under anoxic conditions by utilizing two inflammation-associated metabolites: 1) L-lactate, a non-fermentable carbon source, and 2) nitrate, an anaerobic electron acceptor (44–47). Pathogenic *Neisseria*, such as *N. gonorrhoeae*, similarly depend on L-lactate catabolism for successful host colonization (81, 82), likely exploiting its accumulation during inflammation due to heightened glycolysis in activated immune and epithelial cells (44, 45, 91). Although pathogenic *Neisseria* are unable to respire nitrate (NO_3_^-^), they can survive under anoxic conditions by respiring nitrite (NO_2_ ^-^) (72, 86), a major byproduct (alongside nitrate) of inducible nitric oxide synthase (iNOS) activity (46, 57, 59). While critical for host defense, the upregulation of iNOS during airway inflammation likely creates a niche for nitrate-respiring taxa by increasing local nitrate and nitrite availability (46, 47). These parallels suggest that pathogenic and commensal *Neisseria* capitalize on inflammation-derived metabolites to support growth in oxygen-limited environments.

While lactate and nitrate metabolism may promote *N. mucosa* invasion of the inflamed lower airway, these pathways are most likely maintained because they support the organism’s commensal lifestyle. For instance, we found that nitrate respiration is required for *N. mucosa* to subsist on human saliva, a major nutrient source in the oral cavity (92). This aligns with the known accumulation of dietary nitrate to high (millimolar) levels within saliva, as well as the widespread capacity for nitrate respiration among oral commensals (88, 93, 94). Additionally, lactate is a major fermentation product of commensal oral streptococci, which “cross-feed” lactate to neighboring lactate-utilizing taxa, fostering physical co-association between producers and consumers (15, 89). Thus, the ability to utilize lactate and nitrate may have evolved primarily to support *N. mucosa* colonization of the healthy upper airway, but is co-opted to support its survival in the inflamed lower airway.

This metabolic conservation underscores a key point: while pathogens likely retain nitrate and lactate utilization as adaptations to inflammation, commensals like *N. mucosa* may have developed these functions in the absence of inflammation, but benefit opportunistically when such metabolites are abundant during disease. Indeed, other oral commensals such as *Veillonella*, which also metabolize lactate and nitrate, have been observed to translocate to distal inflamed sites (e.g., the gut of patients with inflammatory bowel disease), raising the possibility that inflammation inadvertently selects for organisms pre-adapted to health-associated metabolic niches (7, 95, 96). In the lung, oral commensals likely reach the lower airways via micro-aspiration, where the metabolic overlap between health and disease contexts facilitates their expansion in chronic infection.

Beyond microbe-host interactions, our results point toward microbe-microbe competition as another critical factor shaping airway colonization. As noted above, nitrate utilization is not unique to *Neisseria*; it is also widespread among other oral taxa, such as *Rothia* species, which co-expand alongside *Neisseria* in response to dietary nitrate and are frequent inhabitants of the cystic fibrosis lung (94, 97). The presence of multiple nitrate-respiring commensals in this niche illustrates how nutrient availability, particularly of alternative electron acceptors like nitrate, can shape microbial community structure (98). As a result, *N. mucosa* is likely to experience niche overlap not only with canonical bronchiectasis pathogens like *P. aeruginosa* but also with fellow nitrate-respiring oral commensals, all competing within the same resource-limited airway environment.

In chronic lung infections, prolonged immune pressure can drive bacteria to evolve reduced immunogenicity, often resulting in loss of inflammatory surface structures such as flagella or pili (99, 100). Interestingly, our Tn-seq data revealed a striking enrichment of pili genes among mutants with enhanced fitness in SCFM2 (**Fig. 2B**), contributing to enrichment of the COG categories “Intracellular trafficking and secretion” and “Cell motility” (**Fig. 2C**). This suggests that pili expression may be disadvantageous for *N. mucosa* under sputum-like conditions, even in the absence of direct host interactions. However, upon further analysis, we found that *N. mucosa* mutants lacking type IV pili machinery also showed increased fitness in standard laboratory media (TSBYE), suggesting this phenotype is likely an artifact of in vitro culture rather than a disease-specific adaptation (**Fig. S2**).

Our study also highlights “phenotypic,” or environmentally induced, antibiotic resistance (65), a phenomenon previously observed among other respiratory pathogens under sputum-like conditions (101, 102). Specifically, we found that *N. mucosa* exhibits increased resistance to multiple, clinically relevant antibiotics when cultured on SCFM2. While the underlying mechanism remains unclear, our Tn-seq screen identified all three subunits of the Mtr efflux pump (**Fig. 2B**, black points), well-known in pathogenic *Neisseria* for conferring antibiotic resistance, particularly to azithromycin (103, 104).

As validation, we constructed a *mtrD* deletion mutant, which, as expected, displayed increased susceptibility to azithromycin and ceftriaxone, though not amoxicillin, on nutrient-rich TSBYE (**Fig. S3A-D**). However, loss of *mtrD* did not affect SCFM2-enhanced ceftriaxone resistance (**Fig. S3D**) and only partially blunted phenotypic resistance to azithromycin. Specifically, while the MIC of azithromycin for the WT increases >60-fold on SCFM2 (**Fig. 1A**), the *ΔmtrD* mutant still showed an 11-fold increase (**Fig. S3A**). These results indicate that the Mtr efflux pump mediates, at least partially, phenotypic antibiotic resistance under sputum-like conditions. Further work is needed to elucidate the complete mechanisms, as well as identify the specific sputum-derived factor(s) that trigger this response.

Given the limitations of conventional antibiotics (50, 63, 64), our findings provide proof-of-concept that targeting infection-specific metabolic pathways may be a promising avenue for the discovery of alternative therapeutics, particularly those overlooked in traditional drug screens using nutrient-rich media. For example, building on pioneering work in the gut microbiome field (60, 105), we found that nitrate respiration—a pathway required for *N. mucosa* fitness in SCFM2 and saliva—can be selectively inhibited using tungstate, a molybdo-enzyme antagonist. While tungstate’s use in the clinic may be limited by potential toxicity (106), its effectiveness against both *N. mucosa* and *P. aeruginosa* in disease-relevant media supports the broader therapeutic potential of targeting nitrate respiration, or other inflammation-associated metabolic dependencies. This approach could be relevant not only for chronic diseases of the lower airway, but also at other barrier sites such as the skin, vaginal tract, or oral cavity.

In conclusion, our study reveals that *N. mucosa* leverages a set of inflammation-compatible metabolic strategies, likely evolved for life in the healthy oral cavity, to persist in the inflamed lower airway. By uncovering the physiological requirements that enable this opportunistic shift, we lay groundwork for the development of novel, non-antibiotic therapeutics that target context-specific metabolic vulnerabilities at the site of infection, with potential application across a range of chronic inflammatory diseases.

## Materials and Methods

### Strains, media, and growth conditions

*Neisseria mucosa* ATCC 19696 and *Pseudomonas aeruginosa* PA14 were used as wild-type (WT) strains. Both were routinely cultured in tryptic soy broth + 0.5% (w/v) yeast extract (TSBYE), or on solid TSBYE + 1.5% (w/v) agar. Synthetic cystic fibrosis medium (SCFM2) (51) was purchased from SynthBiome (#10002, version not zinc-limited). SCFM2 was aliquoted and stored at -20°C until use. A modified version of Socransky’s chemically defined medium (CDM) (107) was prepared as described (108), with CaCl_2_ omitted. Pooled saliva (15-20 mL) was collected from 3-4 healthy human donors, centrifuged to remove debris (∼3,200 x g, 5-10 min), filter-sterilized (0.2 µm pore size), aliquoted, and stored at -20°C until use. Oxic cultures were incubated in 5% CO_2_. Anoxic cultures were incubated in an anaerobic chamber (Coy Laboratory Products or Don Whitley Scientific) containing 90% N_2_, 5% CO_2_, and 5% H_2_. Both oxic and anoxic liquid cultures were incubated without shaking. For some anoxic assays, media and polystyrene culture tubes were pre-reduced overnight in either an anaerobic chamber or, for SCFM2, at 4°C in anaerobic jars (using sachets that generate anoxic conditions). PALA (NSC-224131) (109) was a generous gift from Dr. Christine McDonald (Cleveland Clinic).

### Growth assays

Colonies from streak plates were inoculated into TSBYE and incubated overnight (16-24 h) under oxic conditions. Cultures were pelleted (4,000 x g, 1-2 min), washed with PBS, and adjusted to OD_600_ = 1.0 (or 0.5) in PBS. Cells were then diluted 1:1,000 (or 1:500) into test media to initial OD_600_ = 0.001 (∼10^5^ CFU/mL). Cultures were incubated for ∼24 h under oxic or anoxic conditions before assessing growth, either by measuring OD_600_ or determining CFU (the latter being necessary for SCFM2 due to its high turbidity). For competition assays, WT and mutant strains were both adjusted to OD_600_ = 1.0 (or 0.5), mixed at a 1:1 ratio, and diluted 1:1,000 (or 1:500) into test media. For co-culture with *P. aeruginosa*, *N. mucosa* was adjusted to OD_600_ = 0.5 and *P. aeruginosa* to OD_600_ = 0.005 prior to 1:500 dilution into test media. Growth assays were conducted in 1-mL volumes using 4-mL polystyrene tubes (12 x 75 mm). Where indicated in figure legends, test media were de-oxygenated prior to inoculation. De-oxygenation was not performed routinely until after it was found to be critical for experiments involving tungstate.

### CFU determination

Colony forming units (CFU) were determined as described (110). Briefly, cultures were serially diluted in PBS in 96-well plates (180 µL/well), and using a multichannel pipette, 5 μl of each 10-fold dilution was spotted 5 times onto TSBYE agar (1 plate per sample, 25 μL total per dilution). For *N. mucosa* mutants, dilutions were spotted onto TSBYE agar + 40 µg/mL kanamycin.

### MIC assays

Minimum inhibitory concentrations (MICs) were determined using MIC test strips (Liofilchem) on 0.5x TSBYE and SCFM2 agar. Half-strength agar was prepared by mixing 1x liquid media (either 1x TSBYE + 20 mM NaCl or 1x SCFM2, warmed to 37°C) with an equal volume of autoclaved 2x agar (3% w/v, cooled to 55°C). Lawns were formed by spreading PBS-washed overnight cultures (OD_600_ = 1.0) onto plates using cotton swabs. After a 4-h pre-incubation under oxic conditions (to allow induction of resistance mechanisms), MIC strips were applied, and plates incubated for an additional ∼20 h under oxic conditions. Anoxic MICs were attempted but not reported due to poor lawn formation, even on 0.5x TSBYE agar + 10 mM NaNO_3_.

### *N. mucosa* Tn mutant pool

The *N. mucosa* transposon (Tn) mutant pool was generated by adapting a previously described method for *Acinetobacter baylyi* (111). In this approach, genomic DNA is mutagenized in vitro using EZ-Tn5 transposase (Biosearch Technologies) and then introduced into the target organism via natural transformation. The EZ-Tn5 transposon was generated by PCR-amplifying the kanamycin resistance cassette from plasmid pYGK (112) using 5’-phosphorylated primers containing EZ-Tn5 transposase recognition sequences at their 5’ ends (see **Table S1** for sequences). The in vitro mutagenesis and gap-repair reactions were performed as described in the original protocol but scaled up 5-fold: 8 µg of *N. mucosa* genomic DNA was mutagenized in a 150-µL reaction and gap-repaired in a 250-µL reaction. Parallel negative control reactions substituted H_2_O for both transposase and transposon.

To perform the natural transformation, a colony of *N. mucosa* was re-struck onto TSBYE agar and incubated overnight under oxic conditions. Cells were harvested from the plate into TSBYE + 5 mM MgCl_2_, adjusted to OD_600_ = 2.0, and mixed 1:1 with the unpurified 250-µL gap repair reaction. The mixture was spotted onto 5 polycarbonate filters (0.2 µm pore size) placed on TSBYE agar and incubated for 2.5 h under oxic conditions. Cells were then harvested from the filters by vortexing into ∼10 mL TSBYE + 25% glycerol and stored as 2-mL aliquots at -80°C. One aliquot was thawed, diluted in TSBYE, and spread across ∼30 150-mm TSBYE agar plates + 40 µg/mL kanamycin, followed by incubation for ∼36 h under oxic conditions. The resulting transformants (on average, >2,000 colonies per plate) were harvested into ∼100 mL TSBYE + 25% glycerol and stored as 1-mL aliquots at -80°C. No transformants were observed for the negative-control gap repair reaction.

### Tn-seq experiments

Tn-seq was performed on the *N. mucosa* Tn mutant pool under 4 conditions: input, SCFM2, nitrate, and oxygen. Each condition was tested in biological triplicate (on separate days), for a total of 12 samples. At the end of each experiment, cultures were pelleted and stored at -20°C until further processing.

Input condition: For each replicate, a frozen aliquot of the *N. mucosa* Tn mutant pool was thawed, and 0.5 mL was inoculated into 50 mL TSBYE in a 250-mL flask. Cultures were incubated under oxic conditions for 17-20 h, corresponding to 6-7 generations based on initial vs. final CFU/mL.

SCFM2 condition: Each replicate was initiated from an input culture as described above. A portion of the culture was pelleted at 4,000 x g for 1 min, resuspended in SCFM2, and adjusted to OD_600_ = 1.0. The suspension was then diluted 1:1,000 (initial OD_600_ = 0.001) into 50 mL SCFM2 in a 250-mL flask and incubated under anoxic conditions for 24 h (8-9 generations based on initial vs. final CFU/mL). SCFM2 was not pre-reduced for Tn-seq experiments.

Nitrate and oxygen conditions: Each replicate was initiated from an input culture as described above. A portion of the culture was directly diluted to initial OD_600_ = 0.01 (i.e., without intermediate OD adjustment) in 50 mL de-oxygenated TSBYE + 10 mM NaNO_3_ (nitrate condition) or 10 mM NaCl (oxygen condition) in a 250-mL flask, and incubated under anoxic or oxic conditions, respectively, for 24 h (7-8 generations based on initial vs. final OD_600_).

### Tn-seq library preparation

Genomic DNA was isolated from cell pellets as described (113), with minor modifications (specifically, mutanolysin and lyticase were omitted from the enzymatic lysis step). Tn-seq libraries were then prepared using a two-step PCR method, largely as described (114). Briefly, genomic DNA was: 1) sheared, 2) C-tailed using terminal deoxynucleotidyl transferase (TdT), 3) PCR-amplified for 15 cycles using a 5’-biotinylated Tn-specific forward primer and a C-tail-specific reverse primer (olj376), 4) purified using streptavidin-coated magnetic beads, and 5) PCR-amplified for 20 additional cycles using a nested Tn-specific forward primer and an olj376-specific reverse primer, both containing Illumina adapter sequences at their 5’ ends. Pooled libraries were sequenced on an Illumina NovaSeq (2×150 reads) at the Genome Technology Access Center (McDonnell Genome Institute, Washington University in St. Louis).

### Tn-seq analysis

Tn-seq data were analyzed largely as described (114), using only the R1 read from paired-end sequencing. The analysis proceeded in five main steps.

1) Read pre-processing (Cutadapt v4.6): a) Reads were filtered for and trimmed of the 5’ Tn sequence TTCAGATGTGTATAAGAGACAG, b) trimmed to remove 3’ C-tails (≥12 consecutive C’s) and low-quality bases (Q < 20, fastq format), and c) truncated to a length of 20-25 bp.
2) Alignment (Bowtie 2 v2.5.3) (115): Processed reads were aligned to a re-sequenced assembly of the *N. mucosa* 19696 genome (described below) using default end-to-end settings.
3) Filtering and insertion site analysis (command line): a) Lower-quality alignments (Q < 40, sam format) were discarded, and b) unique Tn insertion sites per sample were identified and counted.
4) Read counting (Rsubread v2.12.3) (116): Read counts per gene (i.e., Tn mutant abundance) were quantified using featureCounts() with a saf annotation file derived from the re-sequenced genome.
5) Differential analysis (DESeq2 v1.42.1) (117): A single counts matrix including all 12 samples was used to normalize read counts for library size and to perform differential abundance analysis using the Wald test with Benjamini-Hochberg correction for multiple testing.

A summary of each analysis step is provided in **Dataset S1**, along with the following files: the saf-format annotation file used with Rsubread, the DESeq2-normalized counts matrix, and the DESeq2 results files for the comparisons SCFM2 vs. input, nitrate vs. input, oxygen vs. input, and nitrate vs. oxygen.

### *N. mucosa* genome re-sequencing

The *N. mucosa* 19696 genome was re-sequenced to support accurate Tn-seq analysis, as the existing NCBI assembly (ASM302831v1, as of 6/18/25) was flagged as contaminated. Genomic DNA was submitted to SeqCenter for hybrid Nanopore long-read / Illumina short-read sequencing, de novo assembly, and genome annotation. The resulting gff annotation file was modified in Excel to create the saf-format annotation file for Rsubread. KEGG K-number annotations were generated by: 1) extracting all gene nucleotide sequences from the re-sequenced genome using the BEDTools getfasta function (118), and 2) submitting the resulting fasta file to the KEGG Automatic Annotation Server (119) using the following settings: BLAST as the search algorithm, the prokaryote representative gene set, and the BBH (bi-directional best hit) assignment method. COG annotations were generated as described (120).

### *N. mucosa* deletion mutants

Deletion mutants were generated by naturally transforming *N. mucosa* with DNA constructs designed to replace the target gene with a kanamycin resistance cassette. Each construct consisted of the kanamycin cassette flanked by ∼1-kb regions upstream and downstream of the target gene. To assemble constructs, the flanking regions were PCR-amplified from *N. mucosa* 19696 genomic DNA, and the kanamycin cassette from plasmid pYGK (112) (see **Table S1** for primer sequences). Assembly was performed using the HiFi DNA Assembly Master Mix (New England Biolabs). Primers for the flanking regions were designed using the NCBI *N. mucosa* 19696 genome assembly (ASM302831v1), and included specific overhangs to facilitate both transformation and assembly: 1) The upstream forward and downstream reverse primers included the *Neisseria* DNA uptake sequence (DUS) at their 5’ ends to enhance natural transformation (121). 2) The upstream reverse and downstream forward primers included sequences complementary to the ends of the kanamycin cassette to enable DNA assembly. Full-length assembled constructs (∼3 kb) were PCR-amplified using the upstream forward and downstream reverse primers, and were confirmed by agarose gel electrophoresis. For transformations, *N. mucosa* colonies were re-struck onto TSBYE agar and incubated overnight under oxic conditions. Cells were then harvested into TSBYE + 5 mM MgCl_2_, adjusted to OD_600_ = 2.0, and mixed 1:1 with the DNA construct (≥500 ng in 50 µL). The mixture (100 µL total) was spotted onto a polycarbonate filter (0.2 µm pore size) placed on TSBYE agar and incubated for 2.5 h under oxic conditions. Cells were subsequently harvested into TSBYE and plated on TSBYE agar + 40 µg/mL kanamycin. Transformants were screened by PCR to confirm deletion of the target gene.

### Statistical analysis

Two-tailed Mann-Whitney tests were performed using GraphPad Prism. Wald tests for Tn-seq data were conducted in R using the DESeq2 package with default settings. One-sided hypergeometric tests were performed in Microsoft Excel using the hypgeomdist function.

## Data Availability

Raw Tn-seq and whole-genome sequencing data have been deposited in the NCBI Sequence Read Archive under BioProject accession numbers PRJNA1280941 and PRJNA1280957, respectively.

## Ethics Statement

This study was approved by the Institutional Review Board of Cleveland Clinic (IRB Protocol #24-118). Saliva samples were collected from healthy adult donors, who were provided with an IRB-approved information sheet describing the study. Use of an information sheet in lieu of written informed consent was approved as part of the minimal-risk study protocol.

## Acknowledgements

This work was supported by NIH grants R00DE031372 (to A.S.) and T32HL155005 (supporting A.M.T.). We thank Dr. Christine McDonald for providing PALA, Drs. Rachel Scheraga and Megan Grund for assistance with in vivo pilot studies, and Dr. Lynn Hajjar for thoughtful feedback on the manuscript. We also thank the Genome Technology Access Center (McDonnell Genome Institute, Washington University School of Medicine) for sequencing Tn-seq libraries. The Center is supported in part by NCI Cancer Center Support Grant P30CA91842. This publication is solely the responsibility of the authors and does not necessarily represent the official views of the NIH.

## Author Contributions

A.M.T.: Investigation, Data curation, Writing – review & editing, Funding acquisition

J.K.: Investigation, Data curation

A.S.: Conceptualization, Methodology, Investigation, Formal analysis, Data curation, Writing – original draft, Writing – review & editing, Funding acquisition

---

**Figure S1.**
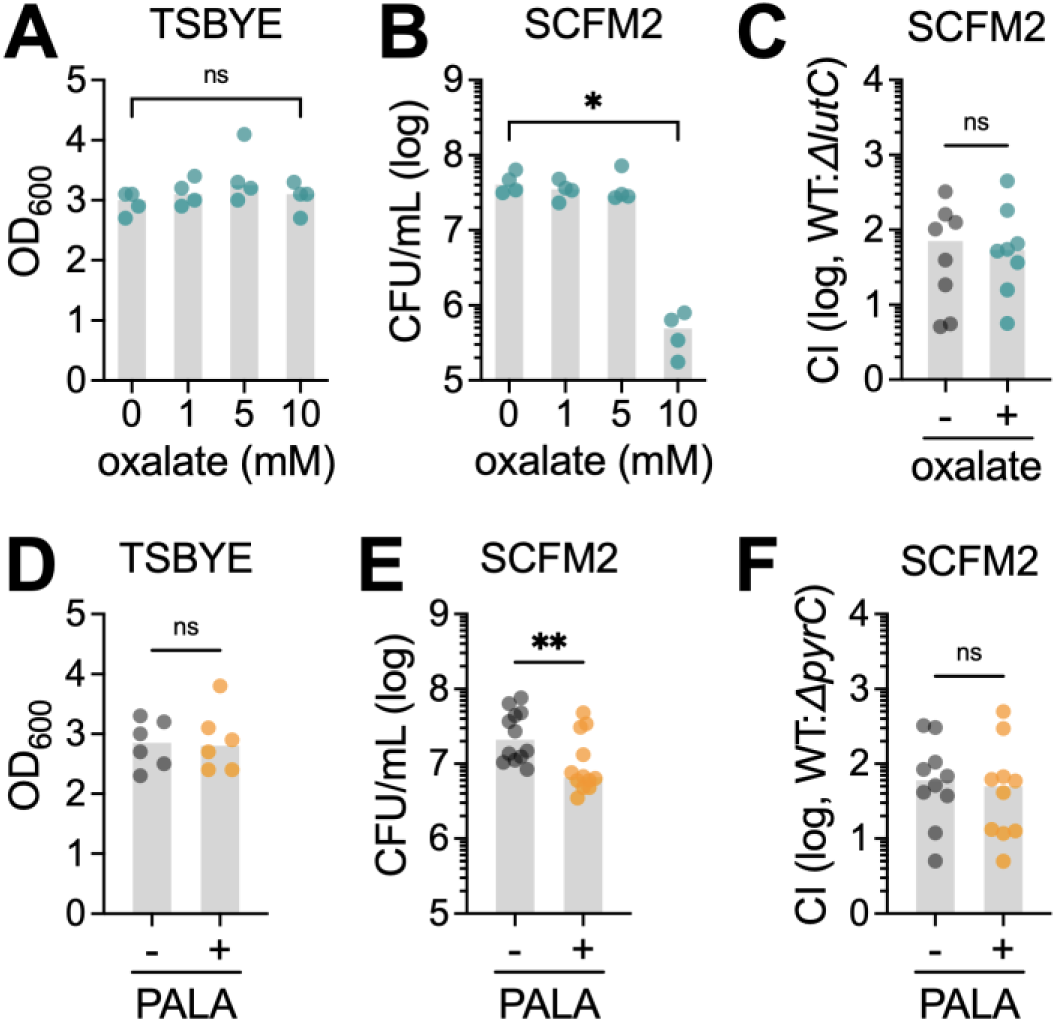
Inhibition of *N. mucosa* L-lactate utilization and pyrimidine biosynthesis in synthetic sputum. (**A, B**) Growth yields of *N. mucosa* after culture for 24 h under anoxic conditions in (A) tryptic soy broth + yeast extract + 10 mM nitrate (TSBYE) or (B) synthetic cystic fibrosis medium (SCFM2) with increasing amounts of oxalate. (**C**) Competitive indexes (CI) for the *N. mucosa* wild type (WT) and L-lactate dehydrogenase mutant (*ΔlutC*) after co-culture for 24 h under anoxic conditions in SCFM2 + 10 mM sodium oxalate or an equal volume of H_2_O (-). (**D, E**) Growth yields of *N. mucosa* after culture for 24 h under anoxic conditions in (D) TSBYE + 10 mM nitrate or (E) SCFM2 + 1 mM PALA (N-phosphonacetyl-L-aspartate) or an equal volume of PBS (-). (**F**) CI for the *N. mucosa* WT and pyrimidine biosynthesis mutant (*ΔpyrC*) after co-culture for 24 h under anoxic conditions in SCFM2 + 1 mM PALA or an equal volume of PBS (-). (C, F) CI determined by dividing the output WT:mutant ratio by the input WT:mutant ratio (∼1:1). (A-F) Media were de-oxygenated in (D-F), but not (A-C), prior to inoculation at OD_600_ = 0.001 (∼10^5^ total CFU/mL). CFU were necessary to assess growth in SCFM2 due to its turbidity. Data represent ≥2 biological replicates (performed on separate days), each with ≥2 technical replicates (total n = 4-12); gray bars, median; *, P < 0.05; **, P < 0.01; ns, not significant (two-tailed Mann-Whitney test).

**Figure S2.**
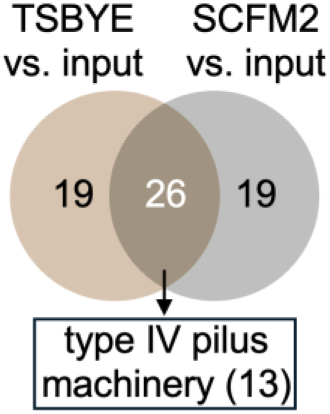
Type IV pili are expendable for *N. mucosa* fitness in vitro. Venn diagram showing the overlap between the number of *N. mucosa* genes in which transposon mutant abundance increased in both tryptic soy broth + yeast extract (TSBYE) and synthetic cystic fibrosis medium (SCFM2) when compared to the input. Of the 26 overlapping genes, 13 coded for components of the type IV pilus machinery.

**Figure S3.**
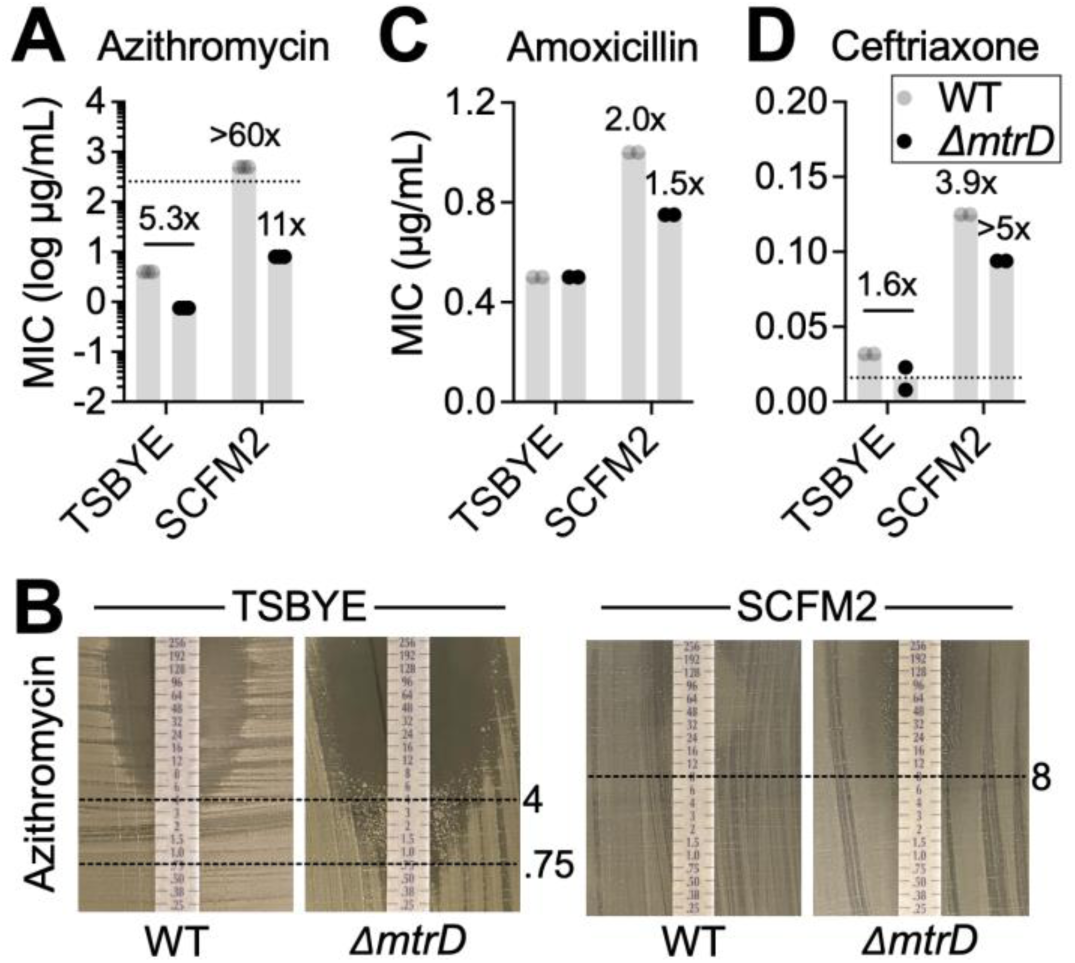
The Mtr efflux pump partially mediates the enhancement of *N. mucosa* antibiotic resistance on SCFM2. (**A, C, D**) Minimum inhibitory concentrations (MICs) of the indicated antibiotics for the *N. mucosa* wild type (WT) and Mtr efflux mutant (*ΔmtrD*) cultured as a lawn on tryptic soy broth + yeast extract (TSBYE) or synthetic cystic fibrosis medium (SCFM2) agar. Lawns were cultured under oxic conditions since growth was not perceptible under anoxic conditions. Numbers above TSBYE bars: fold change in median comparing WT to *ΔmtrD*. Numbers above SCFM2 bars: fold change in median comparing SCFM2 to TSBYE. Dotted lines indicate the max or minimum MIC value. For azithromycin, plotted MIC values for the *ΔmtrD* mutant are the concentrations at which full (not partial) inhibition occurred. (**B**) Representative MIC results for azithromycin. Dotted lines indicate the MIC in µg/mL. Left: top dotted line, MIC for WT; bottom dotted line, MIC for *ΔmtrD*. Right: dotted line, MIC for *ΔmtrD*. (A-D) Data represent ≥2 biological replicates (performed on separate days); gray bars, median.

**Table S1.**
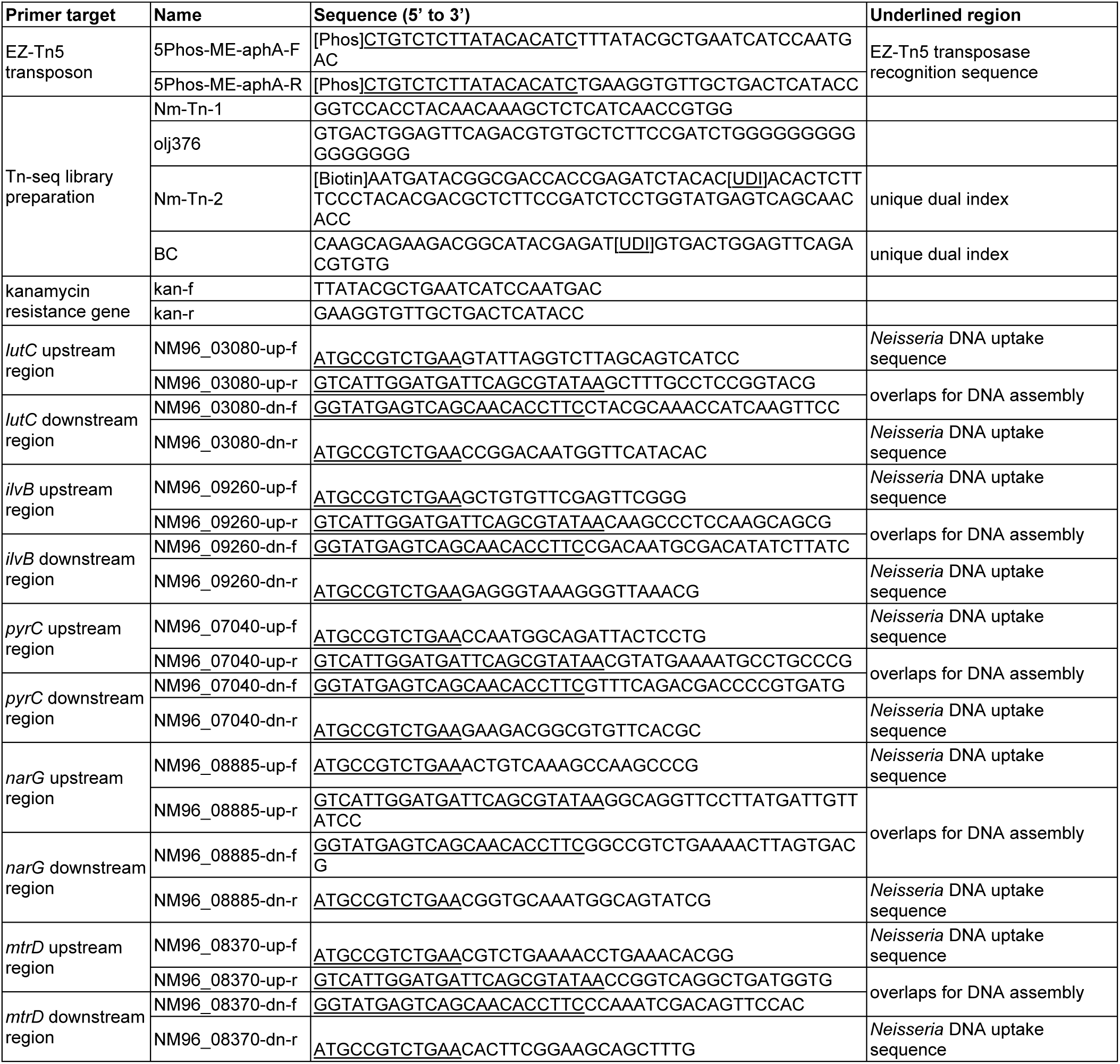
Primers used in this study.

